# EXPOSURE TO GLYPHOSATE OR ROUNDUP^™^ DURING ZEBRAFISH EMBRYOGENESIS DIFFERENTIALLY AFFECTS METABOLISM AND SWIMMING BEHAVIOURS

**DOI:** 10.1101/2022.10.03.510731

**Authors:** Rachel Lacroix, Kingsley Ibhazehiebo, Gaurav Kaushik, Deborah Marie Kurrasch

**Author notes:** These two authors have contributed equally to this work (Rachel Lacroix, Kingsley Ibhazehiebo). To whom correspondence should be addressed: Deborah M Kurrasch, Department of Medical Genetics, Cumming School of Medicine^1^, 3330 Hospital Drive, HSC 2215, Calgary, AB T2N 4N1. **Funding**: This work was funded by NSERC Grant 386445 (DMK).

## Abstract

Glyphosate-based herbicides (GBHs), the most recognized of which is Roundup^™^, are the most extensively used herbicides worldwide. GBHs were initially considered safe for animals since the primary target of the active ingredient, glyphosate, is only found in plants. Recent studies show that glyphosate can affect a range of animal physiologies; however, it remains poorly characterized as to whether animal GBH effects are mediated by glyphosate itself or are in response to toxic effects of the adjuvants and surfactants found in GBH formulations. Here we expose embryonic zebrafish to environmentally relevant doses of glyphosate or Roundup^™^ and compare effects on two systems in the larvae: cellular bioenergetics, since glyphosate can affect mitochondrial function, and behaviour, as a systems-level readout of lasting developmental changes. We showed that exposure of embryonic zebrafish to low levels of glyphosate resulted in decreased mitochondrially-mediated basal respiration and hypoactive swimming behavior in larval stage, whereas exposure to Roundup^™^ in this same treatment paradigm had the opposite effect, causing increased cellular respiration and increased locomotion. In addition, we also explored generational effects of F1 embryonic glyphosate or Roundup^™^ exposure in the F2, and showed that offspring born to zebrafish exposed to low-level glyphosate or Roundup^™^ during embryogenesis both exhibited reduced mitochondrially-mediated basal respiration and altered locomotion. Combined, these data show that embryonic exposure to glyphosate or the full formulation of GBHs caused differential effects on mitochondrial function and behaviours in vertebrates, with potential lasting effects on future generations.

**Figure 1.**
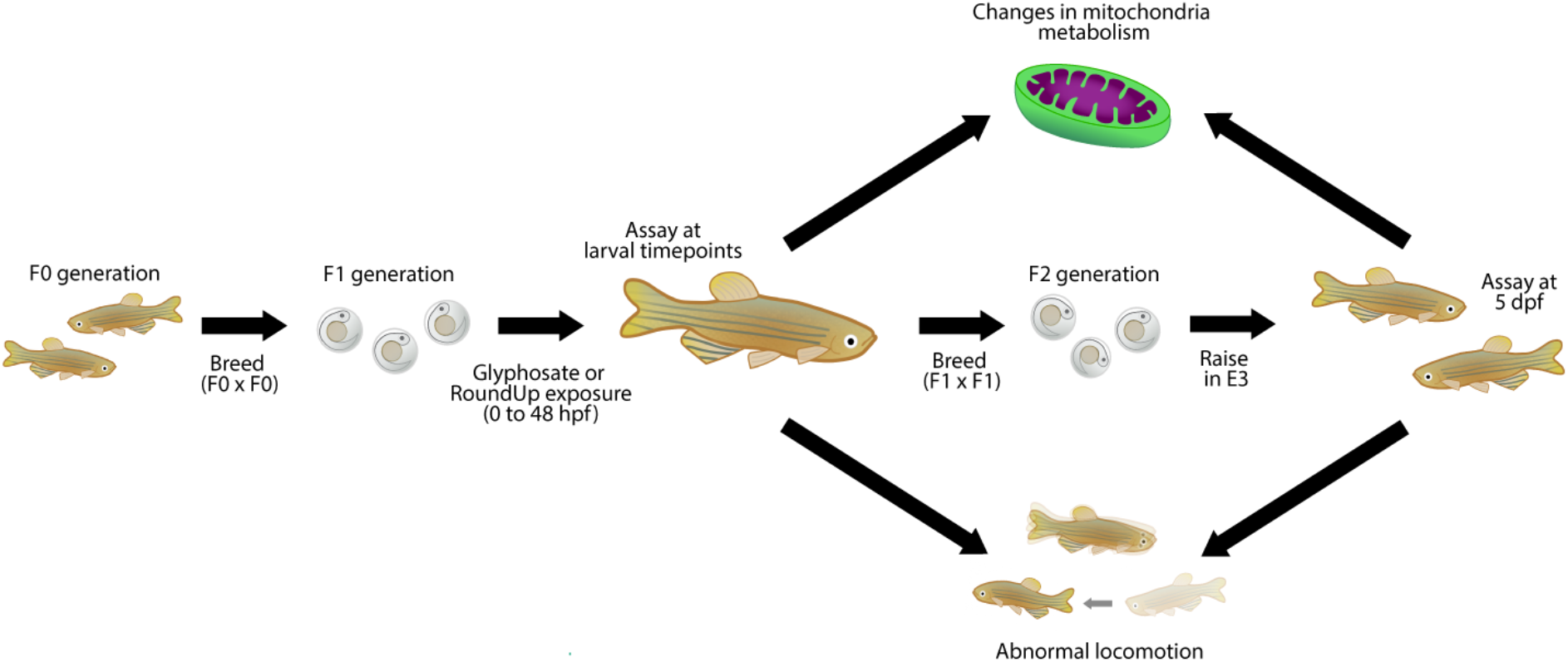
Graphical abstract. Zebrafish (F1 generation) were exposed to glyphosate or Roundup^™^ from zero to 48 hours, with behaviour and mitochondrial bioenergetics assessed at larval timepoints (2- and 5-dpf). Offspring of glyphosate and Roundup^™^ exposed fish (F2 generation) behaviour and mitochondrial bioenergetics were assessed to the same paradigm as F1 fish at 5-dpf. Changes to core metabolic parameters and locomotion was observed in both F1 and F2 generations.

## 1. INTRODUCTON

Glyphosate [N-(phosphonomethyl) glycine] is the active ingredient in glyphosate-based herbicides (GBHs) such as Roundup^™^ and is the most extensively used agricultural herbicide worldwide (Duke and Powles 2008; EPA 2011; Gallardo et al. 2016; Roberts et al. 2010; Wang et al. 2016). GBHs are increasingly used in urban and domestic environments (Botta et al 2009; Brausch and Smith 2007; Webster et al. 2014), furthering animal exposure, including to humans. Agro-based glyphosate usage has reached ~108 million kilograms in the United States and over 800 million kilograms globally since 2014 with sales projected to exceed eight billion USD in 2019 alone (Benbrook 2016; Myers *et al*., 2016). The sharp rise in GBH usage originates in the mid-1990s, when glyphosate-resistant genes were introduced into genetically modified crops (Battaglin *et al*., 2007, 2005). Today, genetically engineered crops with resistance to glyphosate are commonplace; in the United States herbicide-tolerant ‘Roundup Ready®’ soybeans, cotton and corn represent 94%, 95% and 89% of domestic crop acres, respectively (USDA, 2020). Currently, GBHs are applied at various times prior to and throughout the growing season for perennial weed control as well as just prior to harvest to speed crop drying, which results in increased glyphosate and aminomethylphosphonic acid (AMPA; primary metabolite of glyphosate) levels in food sources (Baig 2003; Ellis et al. 1998, Eubank et al. 2008; Giesy et al. 2000; Givens et al. 2009; Bai *et al*. 2016). The increased use of GBHs on Roundup Ready^®^ crops has resulted in 55 herbicide-resistant weed species in over three dozen countries (Heap 2022), propagating a cycle whereby farmers are forced to apply increased levels of herbicides to control these now resistant weeds. Additionally, when GBHs are applied to crops, glyphosate leaches out of the soil, often ending up in surface waters and other aquatic environments (Kanissery *et al*., 2019) that can further reach animal species. In fact, concentrations of glyphosate as high as 5.4 mg/L have been reported in surface waters worldwide, although environmental concentrations are typically in the μg/L range (Solomon *et al*., 2003; Matozzo *et al*., 2020).

Mechanistically, glyphosate targets the rate-limiting enzyme of the shikimate pathway, 5-enolpyruvylshikimate-3-phosphate synthase, which plants require to synthesize essential aromatic amino acids (Hermann and Weaver, 1999). Although glyphosate is marketed as non-toxic to humans and animals due to the absence of this shikimate pathway, emerging evidence indeed suggests negative effects in these non-target organisms upon exposure. For example, glyphosate can disrupt aromatase, a key enzyme in steroid synthesis (Richard et al. 2005). Moreover, glyphosate exposure alone or as a mixture has been evidenced to disrupt mitochondrial metabolism and provokes oxidative damage, as well as activate oxidative stress and compensatory cellular stress response pathways in rats and fish, respectively (Larsen *et al*., 2012; Olorunsogo *et al*., 1979; Webster and Santos, 2015; El-Shenawy, 2009). Rodents exposed to glyphosate display increased risk of developing renal tubule carcinoma, haemangiosarcoma, pancreatic islet cell adenoma, and/or skin tumors (George et al. 2010) and human cell lines treated with glyphosate (50 mg/L) undergo apoptosis (Clair *et al*., 2012; Gasnier *et al*., 2009; Kim *et al*., 2013). Furthermore, another study determined that zebrafish exposed to glyphosate from 1.5 to 120 hpf had transcriptomic-level dysregulation upon low-dose (100 ng/L) exposure to the chemical (Forner-Piquer *et al*. 2021).

The exact formulation of GBHs is proprietary, although it is known that glyphosate is combined with surfactants, commonly polyethoxylated tallow amines (POEA), to increase penetration of glyphosate into plant cells (Botta *et al*., 2009; Brausch and Smith, 2007; Williams *et al*., 2000). Previous studies suggest GBH formulations might be more toxic than glyphosate alone (Annett *et al*., 2014; Defarge *et al*., 2016; Mesnage *et al*., 2013; Moore *et al*., 2012), with GBH co-formulants disrupting endocrine and mitochondrial function in human cells grown in culture at lower concentrations than glyphosate. Furthermore, commercial formulations of GBH containing POEA were 700 times more toxic to amphibians and fish than formulations without POEA (Guilherme et al. 2012; Howe et al. 2004; Perkins et al. 2000), possibly due to adjuvant-facilitated penetration of glyphosate into cells or effects of the surfactants themselves (Annett *et al*., 2014; Bonfanti *et al*., 2018). In juvenile *Danio rerio* (zebrafish), POEA exposure induced greater toxic effects (LC_50_-96 h = 5.49 mg/L) than glyphosate and AMPA (LC_50_-96 h > 100 mg/L); whereas glyphosate induced the highest levels of DNA damage *in vivo* of these compounds (de Brito Rodrigues *et al*., 2019).

Here, we tested environmentally relevant doses (Zhang *et al*., 2016; Matozzo *et al*., 2020) of glyphosate or Roundup^™^, using metabolism and behavior as a readout for cellular and lasting neurodevelopmental effects. We also explored multi-generational consequences of short-term exposure during embryogenesis to glyphosate or Roundup^™^.

## 2. MATERIALS AND METHODS

### 2.1. Zebrafish husbandry and embryo rearing

All protocols and procedures were approved by the Health Science Animal Care Committee (protocol number AC14-0223) of the University of Calgary in compliance with the Guidelines of the Canadian Council of Animal Care. Adult wild type zebrafish (TL strain) were maintained at 28°C in a 14h:10h light-dark cycle in a self-regulating aquatic system (Tecniplast, Buguggiate VA, Italy) with pH 7.5-8.0 and a water conductivity of 980-1000 μS/cm, following a modification of the protocol described by Westerfield *et al*., 2000. The animals were fed twice daily with artemia. The evening prior to the day of breeding, zebrafish breeding pairs were placed in partitioned breeding tank (Tecniplast, Buguggiate VA, Italy) to prevent egg predation. The following morning, spawning was triggered by the onset of light and removal of the partition. Zebrafish embryos were exposed immediately after collection at a density of ~100 embryos/petri dish to approximately equivalent ranges of glyphosate (1.69 μg/L to 1.69 mg/L) or Roundup^™^ (7 μg/L to 7 mg/L acid equivalent (a.e.) of glyphosate; or 10^6^- to 10^3^-fold dilutions) diluted in embryo media as a vehicle (E3; see section 2.3) and were maintained in a non-CO_2_ incubator (VWR, Mississauga, Ontario) at 28°C on a 14h:10h light-dark cycle. Zebrafish embryos did not survive after acute exposure (48 hours) to ≥10^3^-fold dilutions of Roundup^™^, thus only three dilutions that aligned with the lower concentrations of glyphosate used above were tested. These concentrations were chosen to encompass both glyphosate levels in aquatic environments worldwide (~10 ng/L to 10 mg/L; Matozzo *et al*., 2020) and legislated maximum acceptable concentrations of glyphosate in Canadian, American, and European drinking water (0.1 μg/L to 700 μg/L; Health Canada 2005; EU 1998; US EPA 2019). In the metabolic studies, zebrafish were exposed to vehicle (E3), glyphosate or Roundup^™^ from 0-48 hpf, followed by immediate metabolic testing. In the behavior studies, zebrafish were exposed to vehicle, glyphosate or Roundup^™^ from 0-5 dpf, followed by immediate behavioural testing. In the offspring studies, zebrafish embryos were exposed to vehicle, 15 nM glyphosate or a 10^6^-fold dilution of Roundup^™^ for five days, followed by a complete washout of the glyphosate and Roundup^™^ solutions. The glyphosate and Roundup^™^-exposed zebrafish were then grown to adulthood and bred. Embryos of glyphosate- and Roundup^™^-exposed fish (F2 generation) were maintained as previously described in vehicle, and assayed at 5 dpf.

### 2.2. Chemicals

Glyphosate, antimycin A, and carbonyl cyanide 4-(trifluoromethoxy) phenylhydrazone (FCCP) were purchased from Sigma-Aldrich Co (Oakville, Ontario, Canada). Oligomycin was purchased from Enzo Life Sciences (Brockville, Ontario, Canada). Although dozens of RoundUp^™^ formulations are available, we purchased from Walmart the domestically-available, non-selective Roundup^™^ Ready-To-Use with FastAct foam [Pest control product (PCP) no. 22807, Pest Management Regulatory Agency] hereafter referred to as Roundup^™^ throughout the paper. The formulation of Roundup^™^ is proprietary, although it is known to contain 0.96% isopropylamine salt of glyphosate by weight (7 g/L).

### 2.3. Drug and chemical dilution

Embryo media prepared from 333 ml 60X E3, 5 mL methylene blue, diluted in 20 L milli Q water was used as negative control in behavioural and metabolic assays. 100 mM glyphosate was prepared in embryo media and stocked at −20°C. Stock solution of glyphosate was diluted to a final concentration of 1.69 μg/L to 16.9 mg/L in embryo media immediately before use, with repeated freezing and thawing avoided. 10^6^-10^4^x dilutions of commercial Roundup^™^, corresponding to 7 μg/L to 7 mg/L a.e. of glyphosate, was calculated to approximately align with the range of glyphosate concentrations chosen above and was prepared fresh in embryo media immediately before use. 10 mM Oligomycin, Antimycin A and FCCP were diluted in dimethyl sulfoxide (DMSO) or ethanol and stored as a stock at −20°C and diluted to working concentrations immediately before use in the bioenergetics assay.

### 2.4. Metabolic measurements

Oxygen consumption rate measurements of 2-dpf exposed to glyphosate or Roundup for 48 hours and 5-dpf zebrafish exposed to glyphosate or Roundup indirectly through parental exposure were collected using the XF24 Extracellular Flux Analyzer (Seahorse Biosciences, Billerica, MA). Single zebrafish (n = 6-7 per group) were placed in 20 of the 24 wells on an islet microplate and islet plate capture screen was placed over the measurement area to maintain the larvae in a central location. Seven measurements were recorded to establish basal rates, followed by treatment injections of oligomycin, FCCP and Antimycin A and eighteen additional cycles(Stackley et al. 2011). Rates were determined as the mean of two measurements, each lasting two minutes, with three minutes between each measurement (Stackley et al, 2011).

### 2.5. Behaviour assays

5-day post fertilization (dpf) larval zebrafish (n = 12-24 per treatment group) maintained in 96-well plates were habituated for 20 min under ambient light and assayed for locomotor activity and total distance moved in light, dark, and flashing light conditions for 30 min using the Zebrabox (Viewpoint Lifesciences, Lyon, Fr) monitoring equipment at 28°C. All behaviour assays were repeated at least thrice. Total swim activity and tracking of total distance moved as a measure of hyperactive swimming behavior were analyzed using Zebralab V3 software (Viewpoint Lifesciences, Lyon, Fr).

### 2.6. Statistical analyses

Treatment effects were analyzed using one-way ANOVA with post-hoc unpaired t-test or Mann-Whitney test and *p*-values <0.05 were considered statistically significant compared to control.

## 3. RESULTS

### 3.1. Low dose glyphosate decreases respiratory bioenergetics in embryonic zebrafish

We characterized the metabolic profile of zebrafish larvae (2 dpf) directly following exposure to glyphosate from 0-48 hpf by non-invasively measuring bioenergetic changes in real-time. Environmentally relevant doses of glyphosate significantly reduced several measured core metabolic parameters (Figure 2). Specifically, acute (48 hours) exposure to 0.01 and 0.1 μM glyphosate reduced basal respiration (Figure 2a, p<0.01), maximum respiratory capacity (Figure 2b, p<0.001), total mitochondria respiration (Figure 2d, p<0.001), adenosine triphosphate (ATP)-linked respiration (Figure 2e, p<0.05), and respiration due to proton leaks (Figure 2f, p<0.001) in 2 dpf zebrafish. Treatment with 50 μM and 100 μM glyphosate (0-48 hpf) caused similar decreases in basal respiration (Figure 2a, p<0.01), maximum respiratory capacity (Figure 2b, 50 μM only, p<0.05), total mitochondria respiration (Figure 2d, p<0.001), and ATP-linked respiration (Figure 2e, p<0.01) in 2 dpf zebrafish. Notably, exposure to 1 μM glyphosate only affected maximum respiratory capacity (Figure 2b, p<0.05) and exposure to 10 μM glyphosate had no effect on any measured outputs (Figure 2a-f). Non-mitochondria respiration was unchanged in all tested glyphosate treatments (Figure 2c). These data suggest environmentally relevant doses of glyphosate have significant impact on zebrafish bioenergetics during the first few days of life.

**Figure 2.**
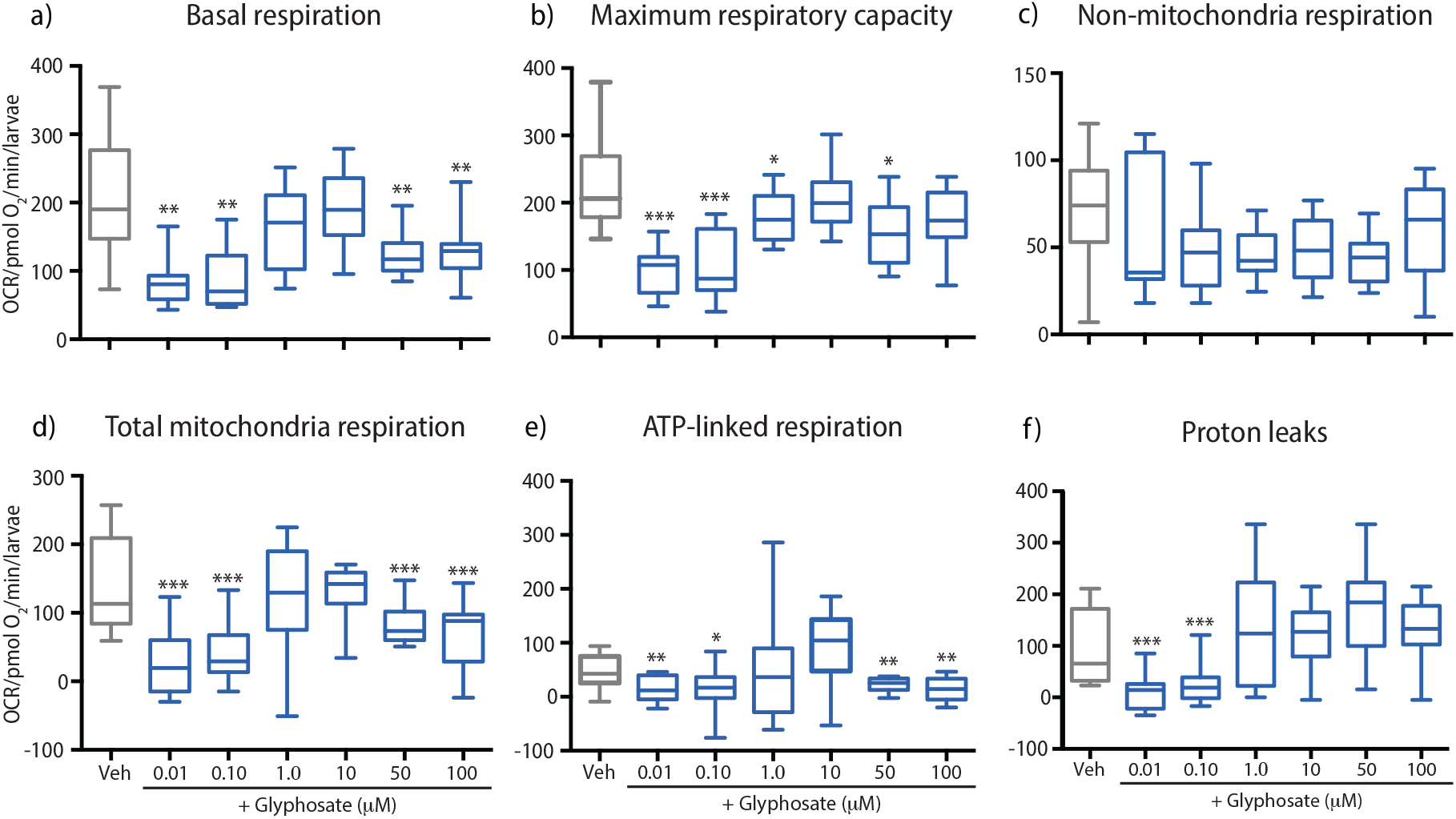
Glyphosate exposure significantly decreases mitochondria bioenergetics in larval zebrafish. (a) Basal respiration; (b) maximum respiratory capacity; (c) non-mitochondria respiration; (d) total mitochondria respiration; (e) ATP-linked respiration; and (f) proton leaks quantified in 2-dpf zebrafish treated following glyphosate (0.01 to 100 μM) treatment. n = 6-7 for all groups. Data are mean ± SE of experiments performed in triplicate. *p<0.05, **p<0.01, ***p<0.001.

### 3.2. Glyphosate exposure during development affects swimming behavior in zebrafish larvae

Given that acute early-life exposure to low-dose glyphosate (0-48 hpf) caused mitochondrial changes in 2 dpf zebrafish, we next investigated if exposure to glyphosate from 0 to 5 dpf caused lasting effects on behaviour of 5 dpf larvae by quantifying movement (pixel movement; “actineg units”; AU) in the dark. We chose this timepoint as the swim bladder has inflated and locomotive behaviours begin to develop. Larvae exposed to 0.01 μM and 0.1 μM glyphosate had significantly decreased locomotion at 5 dpf compared to control treatment, whereas higher concentrations (1-100 μM) had no significant effects on locomotor activity (Figure 3, p< 0.05).

**Figure 3.**
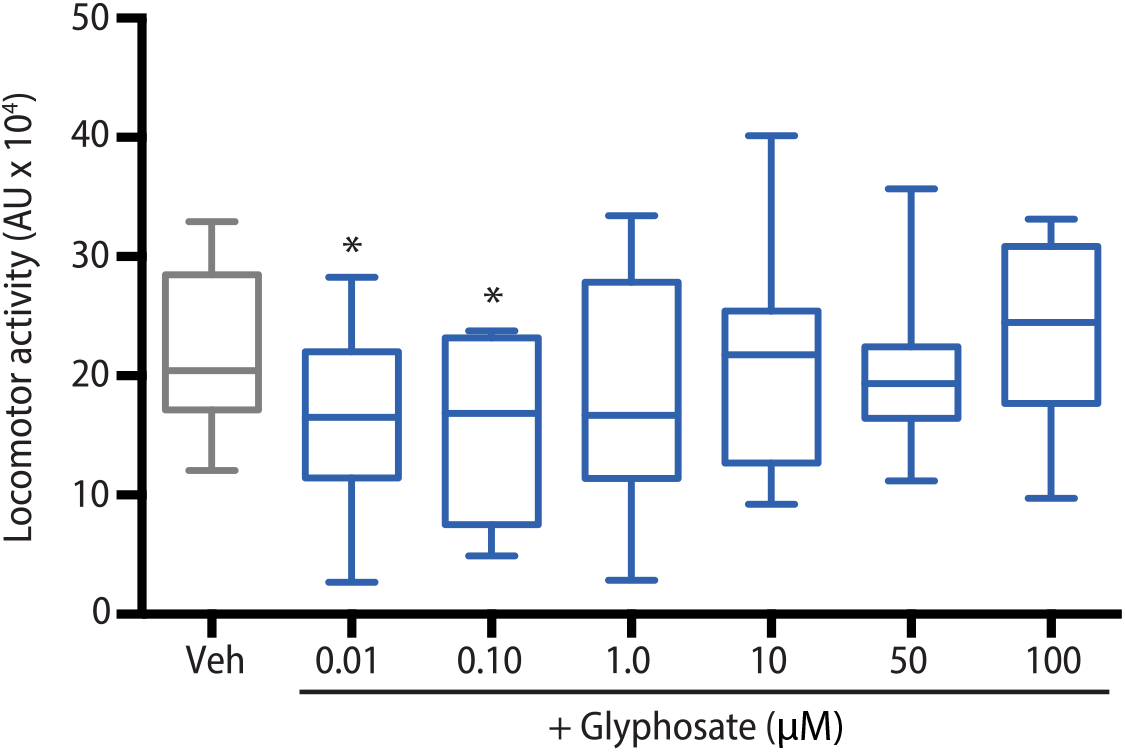
Glyphosate exposure during development causes hypoactive behaviours at low doses. Locomotor activity (actineg unit; AU) of 5-dpf zebrafish exposed to increasing concentrations of glyphosate under dark conditions. N=12-24 for each treatment group. Data are mean ± SE of experiments performed in triplicate. *p<0.05, **p<0.01, ***p<0.001.

### 3.3. Low dose Roundup^™^ increases respiratory bioenergetics in embryonic zebrafish

To test if exposure to a commercial GBH formulation mirrored the changes in zebrafish bioenergetics observed in glyphosate-exposed fish, we exposed embryonic zebrafish (0-48 hpf) to serial dilutions of Roundup^™^. We chose 10^6^-10^4^-fold dilutions of Roundup that approximately aligned with the lower doses of the range of glyphosate doses tested (0.01 μM, 0.1 μM), as these doses caused the most significant effects in the bioenergetics and behaviour assays (Figures 2 and 3), and zebrafish did not survive when exposed to a 10^3^-fold dilution or higher. 48-hour exposure to Roundup^™^ resulted in a significant elevation of several metabolic parameters examined. Specifically, 10^6^-fold dilution Roundup^™^ exposure led to increased basal respiration (Figure 4a, p<0.05), maximum respiratory capacity (Figure 4b, p<0.05), non-mitochondria respiration (Figure 4c, p<0.01), respiration due to ATP turnover (Figure 4e, p<0.01), and significantly reduced respiration due to proton leaks (Figure 4f, p<0.01), where as a 10^4^-fold dilution Roundup^™^ treatment resulted in increased ATP-linked respiration (Figure 4e, p<0.05) and decreased respiration (Figure 4f, p<0.01) due to proton leaks. However, zebrafish embryos exposed to 10^5^-fold dilution Roundup^™^ exhibited decrease in total mitochondrial respiration and respiration due to proton leaks (Figure 4d, 4f, p<0.05).

**Figure 4.**
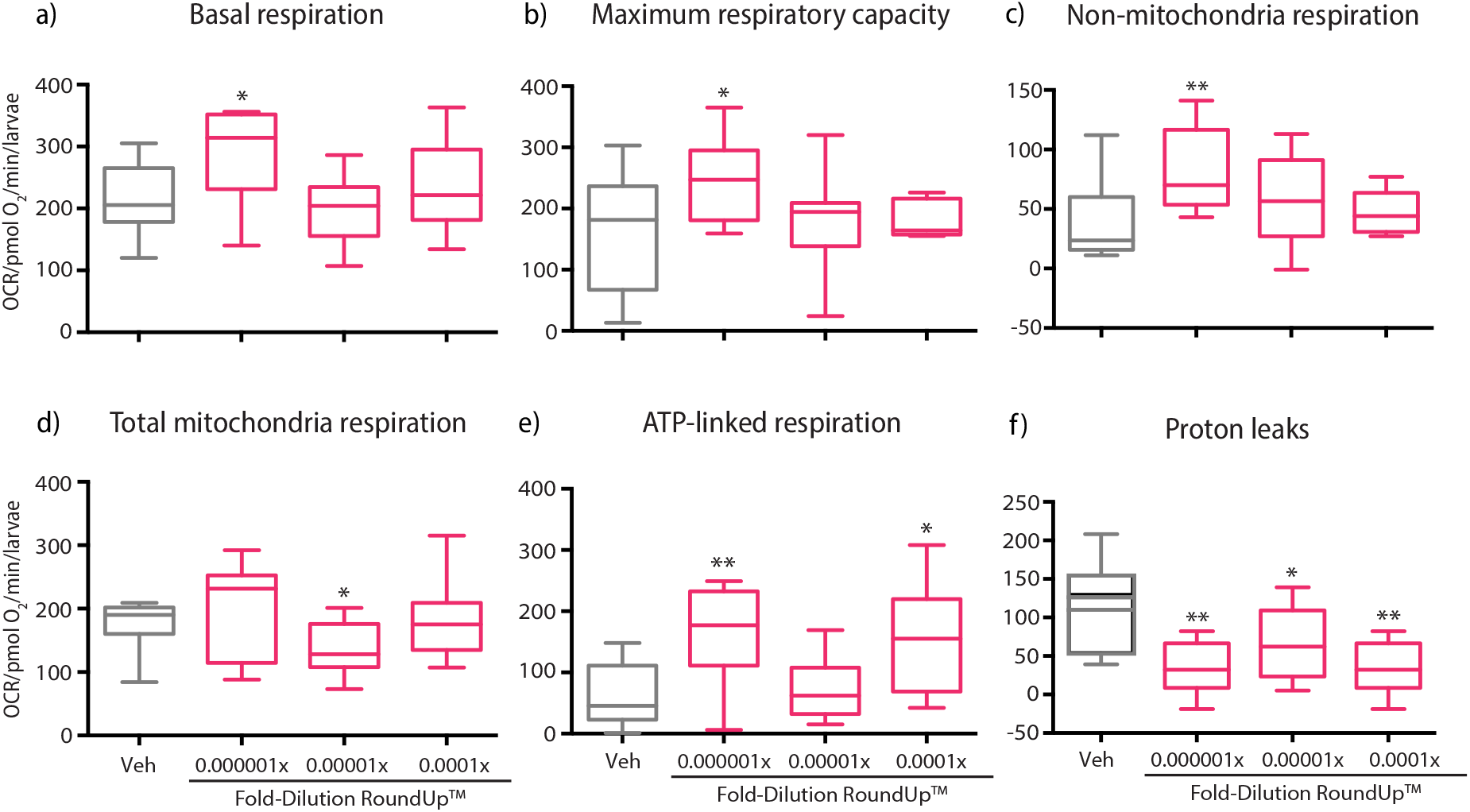
Neurodevelopmental exposure to Roundup^™^ remarkably increases mitochondria bioenergetics in larval zebrafish. (a) Basal respiration; (b) maximum respiratory capacity; (c) non-mitochondria respiration; (d) total mitochondria respiration; (e) ATP-linked respiration; and (f) proton leaks quantified in 2-dpf zebrafish treated with 10^6^ to 10^4^-fold dilutions of commercial glyphosate-based herbicide Roundup^™^. N = 6-7 for all groups. Data are mean ± SE of experiments performed in triplicate. *p<0.05, **p<0.01

### 3.4. Roundup^™^ exposure during development affects swimming behavior in zebrafish larvae

To test if exposure to a commercial GBH formulation mirrored the changes in zebrafish locomotion observed in glyphosate-exposed fish, we exposed zebrafish to serial dilutions of Roundup^™^ as done in the bioenergetics assay. We found that in contrast to glyphosate, zebrafish embryos exposed to 10^6^ to 10^4^-fold dilution Roundup^™^ (0-48 hpf) had significantly elevated locomotor activity from control when assayed at 5 dpf (Figure 5, p<0.05, <0.01, <0.001).

**Figure 5.**
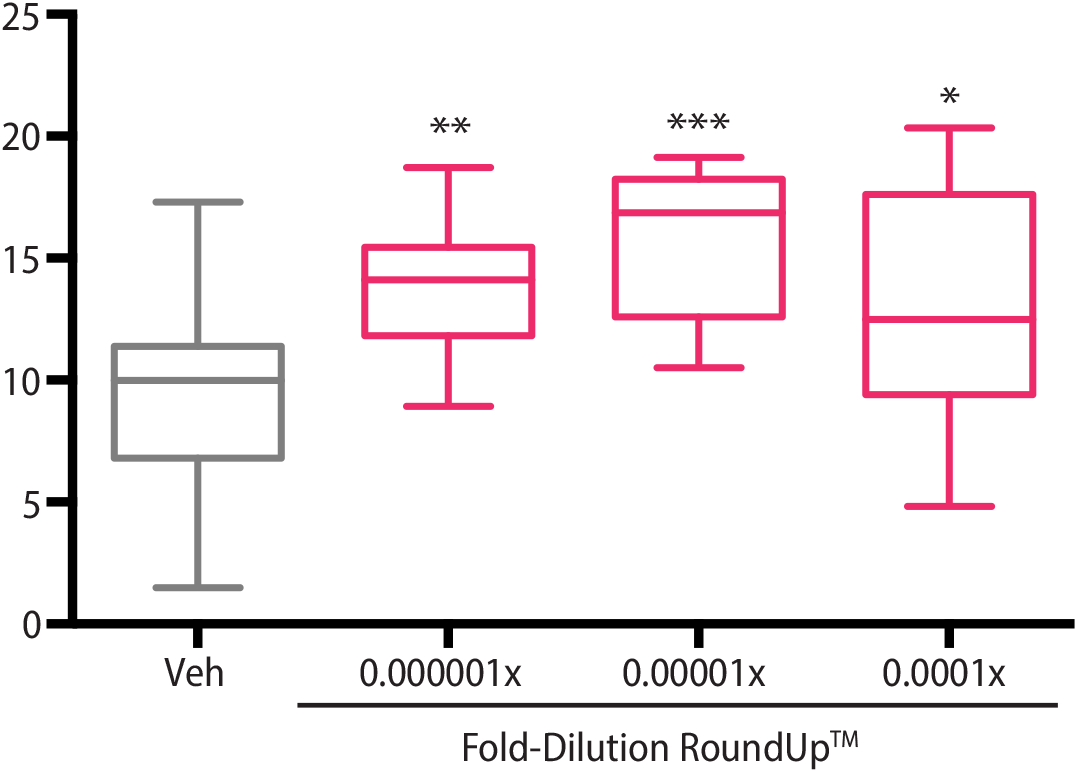
Roundup^™^ exposure during development causes hyperactive swimming across environmental concentrations. Locomotor activity (actineg unit; AU) of 5-dpf zebrafish exposed to Roundup^™^ dilutions under dark conditions. n=12-24 for each treatment group.Data are mean ± SE of experiments performed in triplicate. *p<0.05, **p<0.01, ***p<0.001.

### 3.5. Larval offspring of zebrafish with restricted early-life exposure to low-dose Roundup^™^ and glyphosate have altered bioenergetics and swimming behaviour

Given the changes seen in glyphosate and Roundup^™^-exposed fish, we wanted to investigate if embryonic exposure to glyphosate or Roundup^™^ affects behaviours in offspring of exposed animals. We therefore tested if F1 embryonic exposure to low-dose glyphosate (15 nM) or Roundup^™^ (10^6^-fold dilution) affected F2 larval offspring bioenergetics and/or behaviours. Here, we used 15 nM to capture a dose between low glyphosate doses eliciting significant effects on behavior (Figure 3) that may be relevant to generational studies; Curwin *et al*. determined that the geometric mean glyphosate concentration in the urine of children from non-farming households is 2.5 μg/L (~15 nM) (Curwin *et al*., 2001). F1 zebrafish were exposed to glyphosate or RoundUp^™^ from 0 to 5 dpf, followed by a complete washout until breeding as adults (Figure 6a). We chose a longer exposure period in these studies to encompass primary and secondary neurogenesis, and to capture exposure after zebrafish hatching. 5 dpf F2 offspring of 15 nM glyphosate-exposed parents exhibited decreased basal respiration, decreased total mitochondria respiration, and increased maximum respiratory capacity, with the non-mitochondria component of respiration, ATP-linked respiration, and respiration due protons leaks across the inner mitochondria membrane unchanged (Figure 6b, p<0.05). In contrast to the F1 larvae exposed to RoundUp^™^ that demonstrated increased bioenergetic profiles, the F2 offspring displayed decreases in basal respiration, total mitochondria respiration and respiration due to ATP turnover were remarkably decreased (Figure 6b, p<0.001). Maximum respiratory capacity, non-mitochondria respiration and respiration due to proton leaks were unchanged (Figure 6b). Furthermore, F2 offspring of zebrafish exposed to 15 nM glyphosate displayed significantly hyperactive swimming behaviours when assayed under light conditions (Figure 6c, p<0.05), but there were no significant changes in the dark, as in the case of the F1s who displayed hypoactive swimming under this condition (Figure 6a). F2 offspring of glyphosate exposed F1 zebrafish also had no changes to swimming behaviour when assayed under flashing light. In contrast, F2 offspring of Roundup^™^-exposed F1 zebrafish displayed a decreased in activity when challenged with light-dark flashes but at not in the dark (Figure 6c, p<0.05), as shown for the F1s (Figure 4, p<0.05), or under consistent light (Figure 6c).

**Figure 6.**
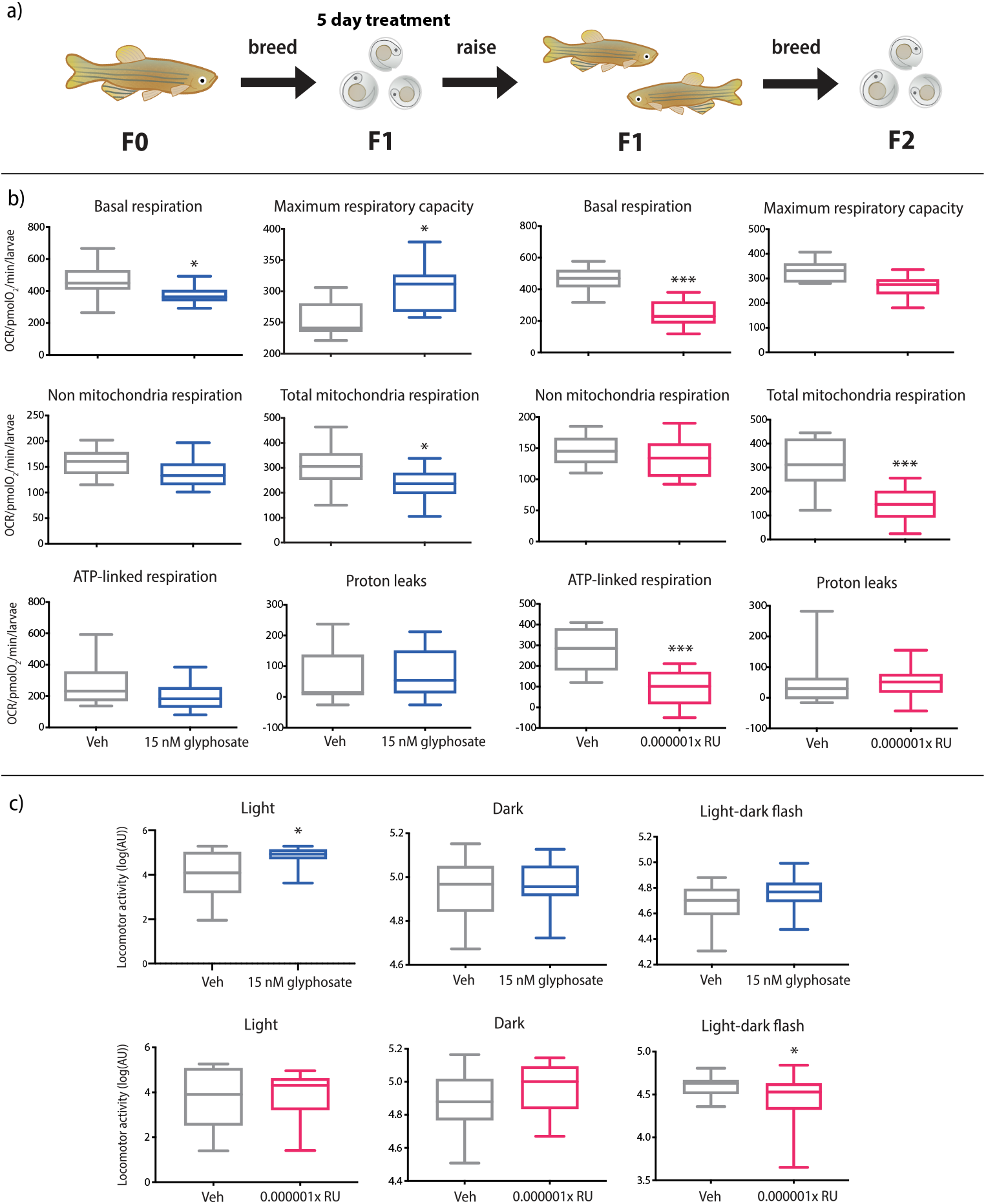
Parental exposure to Roundup^™^ and glyphosate alter bioenergetics and behaviour in larval offspring. (a) Gametes of F1 zebrafish are co-exposed to Roundup^™^ and glyphosate during embryogenesis, which may cause indirect effects in the F2 generation. (b) Quantified basal respiration, maximum respiratory capacity, non-mitochondria respiration, total mitochondria respiration, ATP-linked respiration, and proton leaks in 5-dpf zebrafish treated with 10^6^-fold dilution of commercial Roundup^™^ or 15 nM glyphosate for five days. n = 6-7 for all groups. (c) Locomotor activity [log of actineg unit (AU)] in light, dark or under a light-dark flash (startle assay) in 5-dpf larval offspring following exposure to a 10^6^-fold dilution of Roundup^™^ or 15 nM glyphosate for five days. n = 24 for control and each treatment group. All data are mean ± SE of experiments performed in triplicate. *p<0.05, **p<0.01, ***p<0.001.

## 4. DISCUSSION

In the present study, we report that embryonic exposure to environmentally relevant doses of glyphosate and Roundup^™^ cause changes in mitochondrial metabolism and swimming behaviour in zebrafish. Additionally, offspring of exposed zebrafish show impairment in metabolism and behaviour when tested to a similar paradigm. Our study adds to growing evidence of glyphosate toxicity, showing that embryonic exposure to low-dose glyphosate and Roundup^™^ has bioenergetic and behavioural effects in larval zebrafish, with lasting effects in the subsequent generation.

Despite claims that glyphosate is safe to use, a wide range of toxic effects have been reported across many non-target species (Lopes *et al*., 2014, Dedeke *et al*., 2018, Ruuskanen *et al*., 2020). Our findings add to this growing body of literature by showing that environmental levels of glyphosate and RoundUp affect cellular respiration, with mitochondrial dysfunction likely to impact the overall physiology and metabolism of the animal. Moreover, given that the non-mitochondrial component of cellular respiration remained unchanged in glyphosate-exposed fish, we propose that the toxicity of glyphosate action on bioenergetics is mitochondria specific. In the literature, Panetto and colleagues (2019) used molecular docking experiments in zebrafish to show that glyphosate interacts with the glucokinase, and hexokinases 1 and 2 enzymes involved in respiration, although these effects were seen at a dose closest to our highest tested dose of 100 μM glyphosate, in which changes were observed in this study only on the bioenergetics assay and not in behavior. Interestingly, Pereira and colleagues (2018) determined that zebrafish exposed to 0.065 mg/L of GBH Scout® for 7 days, which is a concentration more similar to our smaller range of glyphosate and Roundup doses, caused behavioural impairments, inhibited complex I and IV activity, and caused mitochondrial hyperpolarization in the brain (Pereira *et al*., 2018). Moreover, adult zebrafish spermatozoa display decreased mitochondrial functionality after short-term glyphosate exposure, though at concentrations exceeding those measured in the environment (5-10 mg/L; Lopes *et al*., 2014). Contrary to our findings, Peixoto *et al*. (2005) report that glyphosate has no effect on mitochondria bioenergetics in isolated rat liver mitochondria. Together, these results suggest glyphosate’s mitochondrial effects are tissue and paradigm dependent, a finding that would be masked by our whole animal *in vivo* approach. Interestingly, in our paradigm mitochondrial bioenergetics have a non-monotonic dose response to glyphosate exposure, with moderate doses (1-10 μM) showing no significant changes across several tested bioenergetic parameters.

Recent studies suggest glyphosate-based herbicide formulations such as Roundup^™^ are significantly more toxic than glyphosate alone (Dedeke *et al*., 2018; Nerozzi *et al*., 2020). The toxicity of GBHs is often attributed to the addition of surfactants, which when applied to plants, break down plant cell walls allowing permeation of glyphosate into cells (Mesnage *et al*., 2019). Polyoxyethylene amines (POEA), surfactants commonly used in GBHs, is thought to contribute to Roundup^™^ genotoxicity more so than glyphosate, with combined glyphosate and POEA having antagonistic effects in fish blood cells when combined (Guilherme *et al*., 2012). In contrast, other studies show that glyphosate and adjuvants have a synergistic effect when administered together in a GBH (Bonfanti *et al*. 2018). Complete Roundup^™^ formulations reveal effects particularly on the mitochondria, including changes to succinate-supported respiration in isolated rat liver mitochondria (Peixoto, 2005), disrupted mitochondria function in soil microorganisms (Nicolas *et al*, 2016), and impaired human sperm mitochondria activity (Anifandis *et al*., 2017). Here, we also found that embryonic exposure to a 10^6^-fold dilution of Roundup^™^ affected many aspects of mitochondrial respiration, further demonstrating that mitochondria are a sensitive target of GBHs. We were surprised that glyphosate-exposed fish displayed a decrease in bioenergetic parameters, as opposed to RoundUp^™^-exposed fish, in which bioenergetic parameters increased at the lowest tested dose, with the exception of proton leaks and total mitochondrial capacity. Specifically, low-dose glyphosate exposure resulted in a global reduction of all examined components of cellular respiration except non-mitochondria respiration. Decreased ATP production and reduction in maximal respiration suggest that glyphosate-exposed zebrafish larvae are incapable of responding to their metabolic demands compared to their wild type siblings. On the other hand, the lowest dose of Roundup^™^ used in our study significantly up-regulated most components of cellular respiration examined except total mitochondria respiration (decreased in the 10^5^-fold dilution treatment group) and respiration due to proton leaks.

In our study, we did not directly compare glyphosate and Roundup doses (as 7 g/L of active ingredient of glyphosate), due to the nature of our exposure paradigm. Adjuvants in Roundup increase the permeability of glyphosate to cells, and as a result, it is possible glyphosate in GBH formulations may accumulate at higher rates in tissues versus glyphosate technical (Bonfanti *et al*.2018; Williams *et al*., 2000). This issue is compounded by the kinetics of glyphosate versus GBH breakdown. In laboratory settings, in water the median half-life of glyphosate is ~5.8 days, whereas in field this number is triple, at ~16.5 days (Covaci, 2014), reflecting climactic differences, but perhaps formulation differences, as well. In Roundup, glyphosate is present in the form of isopropylamine (IPA) salt, increasing the solubility, which may affect tissue uptake in aquatic organisms (Tsai, 2019). To our knowledge there are no studies directly comparing the dissipation of glyphosate technical versus glyphosate in GBH formulation in water in a laboratory environment or E3 medium, however, Klátyik *et al*. determined that glyphosate in Roundup Classic dissipated slower than glyphosate alone (IPA salt) in natural water samples, suggesting POEA surfactants prevent adsorption to solid-liquid surfaces in water (Klátyik *et al*., 2017). In recognizing the difference in dissipation kinetics between glyphosate and GBH formulations and the distinct trends that glyphosate and RoundUp show in our work, in future studies, we plan on doing HPLC analysis to see how glyphosate and GBH concentrations in water correspond to actual exposure in zebrafish larvae, and to directly compare doses based on real glyphosate concentrations in medium and in tissues over the exposure period.

We also show that exposure to low-dose glyphosate or RoundUp^™^ during embryonic development persist as alterations in behaviors. These behavior phenotypes relate well to observed phenotypes in the bioenergetics assays; for glyphosate-exposed fish, several bioenergetics parameters are significantly decreased at lower glyphosate doses (Figure 2), directly corresponding with glyphosate doses that significantly decrease locomotor behaviour in 5-dpf fish (Figure 3). Specifically, our findings that locomotor activity decreased in 5 dpf zebrafish acutely exposed to low-dose glyphosate (0.01 μM, 0.1 μM) is consistent with decreased ambulatory activity in mice following glyphosate exposure (Baier *et al*., 2017), as well as reduced distance traveled and reduced ocular distance in larval zebrafish after a 96-hour glyphosate exposure (Bridi *et al*., 2017). For our Roundup^™^ studies, observed behavior and bioenergetic phenotypes also correlated well, with increased bioenergetic parameters (Figure 4) corresponding to increased locomotion in 5-dpf fish (Figure 5). Interestingly, in both exposure conditions (glyphosate and Roundup^™^), quantification of proton leaks was significantly decreased, opposing the general trend in the Roundup^™^ exposed larvae (Figure 4f). This is peculiar as proton leaks should typically correlate with metabolism and the other facets of bioenergetics tested; although it appears that significant increases in ATP-linked respiration may be compensating for this difference, or vice versa (Figure 4e). Moreover, our findings that locomotor activity increased in larval zebrafish following embryonic exposure to Roundup^™^ is in agreement with Zhang *et al*. (2017) who show that 4 dpf larvae exposed to Roundup^™^ from 0.25 to 48 hpf display increased locomotor activity during the daytime. In contrast, Bridi and colleagues (2017) report that Roundup^™^ reduces locomotor activity in adult zebrafish, but not in larvae; however, they exposed 3 dpf zebrafish larvae to Roundup^™^ for 96 hours and then assayed for changes in locomotion whereas our exposure window was 0-48 hours with behaviours analyzed at 5 dpf. Thus, experimental differences may account for the differences between our findings and those of Bridi *et al*.

To the best of our knowledge, no study has investigated the effects of environmentally relevant concentrations of glyphosate or GBHs on metabolic or behavioural profiles in zebrafish offspring (F2) from exposed parents (F1). Offspring from restricted (5 days) Roundup^™^ and glyphosate exposed F1 zebrafish likewise showed impaired cellular respiration, specifically basal and total mitochondria components. Interestingly, maximum respiratory capacity increased in offspring from glyphosate-exposed parents, and ATP-linked respiration decreased in offspring from Roundup^™^-exposed parents. Unexpectedly, we observed that larval F2 offspring from low-dose glyphosate-exposed F1 zebrafish exhibited increased swimming behaviour in light, while those from Roundup^™^-exposed F1 zebrafish showed decreased locomotion under a startle assay (light-dark flash). One plausible explanation for the “glyphosate phenotype” seen in larval offspring of Roundup^™^-exposed zebrafish is that the glyphosate component of Roundup^™^ may persist for longer than the adjuvants, leading to a more lasting effect on physiology that can persist in the offspring. This may be due to epigenetic changes from parental exposure that are further passed on to the offspring, persisting in changed behaviors. Overall, the mechanism by which Roundup^™^ and glyphosate exposure may alter behavior and respiratory bioenergetics in successive generation is still unknown and may be paradigm and model specific. Further studies are needed to identify possible signaling pathways targeted by Roundup^™^ and glyphosate, whether modes of action are different, and how such pathways may contribute to altered metabolism and behaviors in offspring.

Whether and how the zebrafish bioenergetic and behavioural alterations from glyphosate or RoundUp^™^ translate to humans remains to be elucidated. For example, we exposed fish to glyphosate or Roundup^™^ in E3 medium, thereby bathing developing zebrafish in these chemicals, whereas human contact is likely to occur by ingestion or point of contact. As such, it is not clear whether humans exposed to glyphosate or RoundUp^™^ would display equal changes in mitochondrial function given a more restricted point of contact. Moreover, additional studies are needed to understand how altered bioenergetics in humans, if present, manifest as changes in physiology. That said, the glyphosate and GBH exposure to zebrafish in our study does model embryonic exposure of fish in natural habitats, given the environmental relevance of the concentrations used in our study (ng/L-μg/L acid equivalent) and persistence of glyphosate and GBHs in aquatic habitats through soil leaching (Kanissery *et al*., 2019, Matozzo *et al*., 2020). Thus, our findings do provide further evidence that these chemicals do have effects in aquatic animals despite being billed as ‘safe’ to non-plant species.

In summary, here we show that *in vivo* embryonic exposure to GBHs or its active ingredient glyphosate differentially alters metabolism and behaviors, with lasting consequences on the following generation. Although still controversial to what extent, a growing body of evidence shows that glyphosate and Roundup^™^ can persist in animals and humans with potential toxicity. In fact, conclusions drawn in 2017 by The WHO’s International Agency for Research on Cancer (IARC) working group classified glyphosate as a grade 2A probable human carcinogen, although this ruling is highly criticized. Our data here shed additional insights into the behavioral consequences of embryonic GBH exposure and suggest that glyphosate and Roundup^™^ may affect metabolism and behaviour differently, further illustrating the importance of assaying both the active ingredient and commercial product that have adjuvants added.

## Abbreviations

GBH: glyphosate-based herbicide
hpf: hours-post fertilization
dpf: days-post fertilization
POEA: polyethoxylated tallow amine
AMPA: aminomethylphosphonic acid

## Acknowledgments

We would like to Younghee Ahn for training and support on the Seahorse Bioanalyzer and Alicia Vandenbrink for administrative support.

